# Fruit abundance and moonlight shape sex-specific nocturnal activity and movement in kinkajous (*Potos flavus*)

**DOI:** 10.64898/2026.08.23.746512

**Authors:** Roland W. Kays

**Affiliations:** North Carolina Museum of Natural Sciences, Raleigh, NC, USA; Department of Forestry and Environmental Resources, North Carolina State University, Raleigh, NC, USA

## Abstract

Animals face a fundamental trade-off between food-related competition and predation risk in how they allocate time and behavior. Nocturnal mammals offer a particularly tractable system for testing this trade-off because moonlight creates a natural, quantifiable gradient in both predation risk and the visibility needed for safe movement. We used minute-by-minute focal observations of nine kinkajous (*Potos flavus*; 4 female, 5 male) in Panama to test how fruit abundance and moonlight predicted nocturnal activity budgets (percent time traveling, feeding, resting) and nightly travel distance. Beta-family generalized linear mixed models and a linear mixed model of log travel distance showed that fruit abundance was positively associated with percent time traveling and with nightly travel distance, and negatively associated with percent time feeding: kinkajous traveled more and fed less per hour when fruit was abundant, consistent with movement between many nearby productive trees rather than prolonged feeding at a few. Males traveled less as moonlight increased, while females traveled more. Rainfall had no independent effect. We interpret the sex-reversed moonlight response as evidence that moonlight elevates predation risk for males while facilitating movement for the more food-limited females of this frugivorous carnivore.

## INTRODUCTION

Animals face a fundamental trade-off between the time and energy required to locate and acquire food and the risk of predation incurred while doing so; the balance between these two pressures is a classic organizing framework for understanding time allocation and behavior across many taxa (Terborgh and Janson 1986; van Schaik 1989). On the food-competition side of this trade-off, fruit-eating mammals in tropical forests experience marked temporal variation in the abundance of ripe fruit, and when fruit is scarce, animals are expected to spend more time searching and traveling between sparse or depleted patches and correspondingly less time engaged in sustained feeding at any one patch (Milton 1980; Chapman et al. 1992).

Nocturnal mammals offer an especially good system in which to test the balance between food competition and predation risk, because moonlight varies predictably from night to night and across the lunar cycle, creating a natural, quantifiable gradient in both predation risk (through increased visibility to predators) and the light available for safe, efficient movement (through visibility of the substrate itself). For many small nocturnal mammals, brighter moonlight is associated with reduced activity and increased vigilance, consistent with a predation-risk cost of conspicuousness under illumination – the widely reported “lunar phobia” response (Prugh and Golden 2014; Palmer et al. 2017). But moonlight can also facilitate activity in species for which darkness itself limits movement, either by constraining visual navigation or by increasing the physical risk of movement error, a pattern sometimes called “lunar philia.”

The kinkajou (*Potos flavus*) is a strongly arboreal, largely frugivorous procyonid of Central and South American forests whose natural history and social organization at our study site have been described in detail (Kays and Gittleman 2001; Kays et al. 2000; Kays 2003).

Kinkajous forage individually or in small family-based groups, feed on ripe fruit supplemented by nectar, and defend territories against conspecifics (Kays 1999). This resource-defense ecology predicts that fruit scarcity should increase travel and reduce time spent feeding, a pattern consistent with within-patch feeding competition.

Kinkajous also present an unusual test case for the moonlight-movement question because of their body size. At roughly 1.5-3 kg, the kinkajou is the largest nocturnal mammal in the Neotropical forest canopy. While diurnal primates move by leaping confidently between branches and lianas, a kinkajou moving through the canopy at night relies on continuous four-limbed contact with the substrate and a prehensile tail. A fall from the canopy is a real source of injury risk for an animal of this size and locomotor style, and a kinkajou may be more dependent than a leaping primate on being able to see the substrate it is stepping onto. Thus, rather than assuming moonlight simply increases predation risk and thus suppresses activity, we treat it as an open, testable question whether increased ambient illumination instead facilitates safer, more extensive canopy travel in an animal for which darkness itself may be a meaningful locomotor constraint.

Male and female mammals commonly differ in reproductive investment, with females alone bearing the direct metabolic costs of gestation, lactation, and offspring care across most mammalian species (Clutton-Brock 1991). Kinkajous fit this general pattern: females alone bear these reproductive costs (Kays and Gittleman 2001; Kays 2003). Because of this, we expect the two sexes to respond differently to both fruit scarcity and moonlight. A female that is more energetically constrained by reproduction should be more sensitive to variation in food availability, and may also be more willing to trade increased conspicuousness under moonlight for the movement benefits of better visibility if she is under greater pressure to cover ground and find food. We therefore tested (1) how fruit abundance and moonlight predict the nocturnal activity budget (percent time traveling, feeding, and resting) and nightly travel distance, and (2) whether these relationships differ by sex.

## MATERIALS AND METHODS

### Study Site

The study was conducted in the Limbo research plot (104 ha), within Parque Nacional Soberanía (22,100 ha; 9.15°N, 79.7333°W) along Pipeline Road, Panama. The site is tropical moist forest receiving approximately 2,600 mm of rainfall annually, roughly 90% of which falls during the wet season from late April to mid-December. The forest at the Limbo plot is a mosaic of secondary growth 60-120 years old and remnant patches of forest approximately 400 years old (Kays and Gittleman 2001).

### Animal Capture and Marking

Kinkajous were captured using 50 hoistable Tomahawk live traps set in the forest canopy and understory, following the trap design described by Kays (1999). Trapping over the study period yielded 192 captures across 1,292 trap-nights, resulting in 25 individually identified kinkajous. Ten animals were fitted with radio collars and 15 with collars bearing reflective tags for nocturnal visual identification, following the capture and marking protocols used in companion studies of this population (Kays and Gittleman 2001; Kays et al. 2000).

### Travel Distance

Nightly travel distance was measured directly by the observer: while following the focal animal, the observer pace-counted the distance covered from known trail-marker landmarks and recorded the compass bearing of each straight-line travel segment with a hand compass. The resulting route was reconstructed afterward on a paper trail map to calculate the total distance traveled during the tracked interval. This produced 63 nightly travel-distance records (one per individual per half-night), with tracked intervals ranging from 5.0 to 6.33 hours and total distances ranging from 74 to 2,447 m.

### Behavioral Data Collection

Nine of the marked individuals (4 female: Griz, Maggie, Orange, Patty; 5 male: Bear, Elvis, Ernie, Otis, Pokey) were followed on foot during half-night observation sessions, either dusk to midnight or midnight to dawn, between February and December 1996. An observer maintained continuous visual or radio-telemetric contact with the focal animal and recorded its behavior on a minute-by-minute basis, classifying each minute as traveling, feeding, resting, engaged in social behavior, or unknown. Minute tallies were aggregated to hourly percentages of time in each behavior category for each individual within each session. This protocol produced 380 hourly behavioral records nested within 65 half-night sessions (one session, for the individual Elvis, was an uninterrupted full-night observation rather than a half-night). In addition to distance moved, three primary compositional response variables were analyzed: percent time traveling, percent time feeding, and percent time resting.

### Environmental Covariates

#### Fruit abundance

Fruit availability was assessed by a periodic transect census of fallen ripe fruit along an 11.4-km, 0.5-m-wide transect following the established grid of trails within the Limbo plot, following the census method of Kays (1999). This census recorded all fallen fruit meeting the count threshold along the transect, regardless of species, and is therefore a general measure of fruit availability at the site rather than a measure restricted to species known to be eaten by kinkajous. The census was repeated at roughly 4-week intervals throughout the study period, yielding 65 census dates spanning February through December 1996. On each census date, the running tally of fruiting trees encountered along the fixed transect was converted into a fruit-abundance index (an integer count of fruiting trees or discrete fruit patches meeting the threshold criterion along the route) and a companion mean inter-fruiting-tree distance index. Across the 65 census dates, the fruit-abundance index ranged from 2 (the sparsest census date, with very few fruiting trees encountered along the entire transect) to 49 (the most fruit-abundant date), and the mean inter-fruiting-tree distance ranged from 154 to 1,485 m; low index values with correspondingly large inter-tree distances indicate a period of scarce, widely scattered fruit, while high index values with short inter-tree distances indicate a period when fruiting trees were common and closely spaced along the transect. Each behavioral observation was assigned the fruit-abundance value from the most recent preceding census date.

#### Moonlight

Nighttime illumination was quantified from the position and phase of the moon for the exact site coordinates and the date and time of each observation, following the biologically meaningful moonlight-modeling approach of Smielak (2023), producing a clear-sky relative moonlight index; scaled to the illuminance of an average full moon at zenith, i.e., directly overhead. Because the observation hours in this study never coincided with the moon being simultaneously at full phase and at zenith at the same time, the maximum realized value of this clear-sky index in the dataset is well below the theoretical ceiling of 1.0 (observed maximum = 0.673 among the 380 hourly behavioral records).

Because cloud cover attenuates the moonlight that actually reaches the forest floor, this clear-sky index was combined with hourly cloud-cover fraction (Open-Meteo Historical Weather API; Zippenfenig 2023, drawing on ERA5 reanalysis; Hersbach et al. 2020) to produce a second, cloud-adjusted moonlight measure approximating the illumination actually available on the ground; this cloud-adjusted index reaches an even lower realized maximum (0.178) because cloud attenuation is multiplicative with the clear-sky value.

Both the clear-sky and cloud-adjusted moonlight measures were used as alternative versions of the moonlight covariate in parallel sets of models, allowing us to evaluate whether accounting for realized, ground-level illumination sharpened or altered the moonlight relationships observed under the theoretical clear-sky index.

#### Rainfall and breeding season

Hourly precipitation (Open-Meteo Historical Weather API; Zippenfenig 2023, drawing on ERA5 reanalysis; Hersbach et al. 2020) was included as a covariate to test for an independent effect of rainfall once fruit abundance and moonlight were accounted for. A coarse two-level breeding-season index, distinguishing two parts of the annual reproductive cycle, was included as a fixed effect in all models.

### Statistical Analysis

The three activity-budget proportions (percent time traveling, feeding, and resting) were modeled with beta-family generalized linear mixed models (logit link), the appropriate distribution for a bounded proportional response. Because hourly observations within a session are serially correlated, each model included a first-order autoregressive (AR1) correlation structure over within-session hour index, together with a random intercept for individual to account for repeated sampling of the same animals. Nightly travel distance was log-transformed and modeled with a linear mixed model with a random intercept for individual. Fixed effects in all models were fruit abundance, moonlight (fit separately in clear-sky and cloud-adjusted versions), sex, breeding season, and rainfall; models of percent time traveling, percent time resting, and nightly travel distance additionally included a sex-by-moonlight interaction, and an exploratory model of the presence of social behavior additionally included both sex-by-fruit and sex-by-moonlight interactions. All continuous predictors were standardized (z-scored) prior to model fitting so that fixed-effect estimates are directly comparable as standardized effect sizes across covariates and response variables. Models were fit in R (version 4.3 or later) using the package glmmTMB (Brooks et al. 2017), and residual diagnostics for the primary travel-distance model were evaluated using simulation-based quantile residuals in the package DHARMa (Hartig 2022). Effect sizes are reported as standardized coefficients with 95% Wald confidence intervals; the fitted activity-budget and travel-distance surfaces shown in the figures are back-transformed model predictions holding other covariates at their sample mean or reference level, with 95% confidence bands from the delta-method standard error of the linear predictor.

### Data Availability

The raw data and analytical code for this paper are available on Data Dryad

## RESULTS

Across 380 hourly behavioral records from 65 half-night sessions (9 individuals: 4 female, 5 male), fruit abundance was positively associated with the proportion of time spent traveling and negatively associated with the proportion of time spent feeding, while nightly travel distance (63 records) increased with fruit abundance and showed a significant, sex-reversed relationship with moonlight.

### Activity Budget and Fruit Abundance

Fruit abundance had a significant, positive effect on percent time traveling in the clear-sky moonlight model (standardized β = 0.288, SE = 0.137, 95% CI [0.020, 0.556], *P* = 0.035) and a similar, marginally non-significant effect in the cloud-adjusted model (β = 0.250, SE = 0.140, 95% CI [-0.025, 0.525], *P* = 0.074). The corresponding effect on percent time feeding was negative in both models (clear-sky: β =-0.234, SE = 0.132, *P* = 0.076; cloud-adjusted: β =-0.230, SE = 0.130, *P* = 0.078), and the effect on percent time resting was small and not significant in either model (*P* = 0.459 and 0.698). Predicted activity budgets across the observed range of the fruit-abundance index (Fig. 1) show the expected pattern: as fruit abundance increases, predicted percent time traveling rises from roughly 25-30% to 50-60% of the hour, while predicted percent time feeding declines from roughly 40-50% to 25-30%. Females and males followed broadly parallel trajectories across the fruit gradient, with females showing a significantly higher predicted percent time traveling than males at any given level of fruit abundance (sex main effect, β = -0.442 for males relative to females, *P* = 0.020); the corresponding sex difference in percent time feeding and resting was not statistically significant (*P* = 0.300 and *P* = 0.237, respectively; Fig. 1).

**Figure 1.**
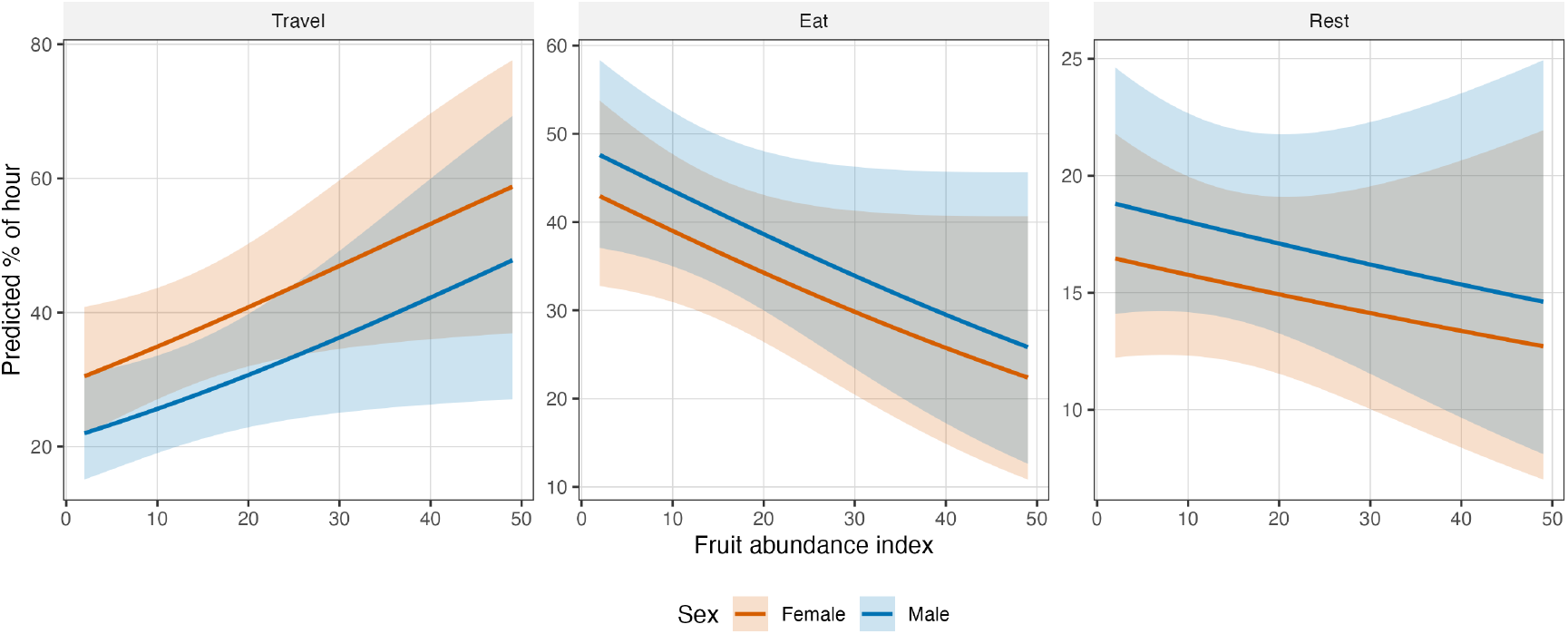
Predicted activity budget (percent time traveling, feeding, and resting per hour) as a function of the fruit-abundance index, separately for females (orange) and males (blue). Lines are model-predicted values from the beta-family generalized linear mixed models (clear-sky-moonlight model version), holding moonlight, rainfall, and breeding season at their sample mean or reference level; shaded bands are 95% confidence intervals. Fruit abundance had a significant positive effect on percent time traveling (β = 0.288, *P* = 0.035) and a non-significant negative effect on percent time feeding (*P* = 0.076) and percent time resting (*P* = 0.459). Females had significantly higher percent time traveling than males across the fruit gradient (sex main effect, *P* = 0.020); the sex difference in percent time feeding and resting was not statistically significant (*P* = 0.300 and *P* = 0.237, respectively). These models did not include a sex-by-fruit-abundance interaction term, so the female and male trend lines are constrained to be parallel across the fruit gradient.

### Activity Budget and Moonlight

Neither the clear-sky nor the cloud-adjusted moonlight index had a significant main effect on percent time traveling, feeding, or resting (all *P* > 0.2). However, the sex-by-moonlight interaction for percent time traveling was significant (*P* = 0.020): predicted percent time traveling increased with moonlight for females (from approximately 42% to 48% of the hour) but decreased for males (from approximately 39% to 22%) across the observed moonlight range (Fig. 2). The parallel sex-by-moonlight interaction for percent time resting was in the same qualitative direction in both moonlight versions and was statistically significant in the cloud-adjusted model (*P* = 0.040), though only marginally non-significant in the clear-sky model (*P* = 0.093): predicted percent time resting rose with moonlight for males (from approximately 14% to 24% under clear-sky moonlight) while remaining comparatively flat for females (Fig. 2). As with nightly travel distance, this interaction was therefore more clearly resolved under the cloud-adjusted, realized-illumination covariate than under the clear-sky index. Percent time feeding did not include a sex-by-moonlight interaction term and showed no significant moonlight or sex effect (Fig. 2).

**Figure 2.**
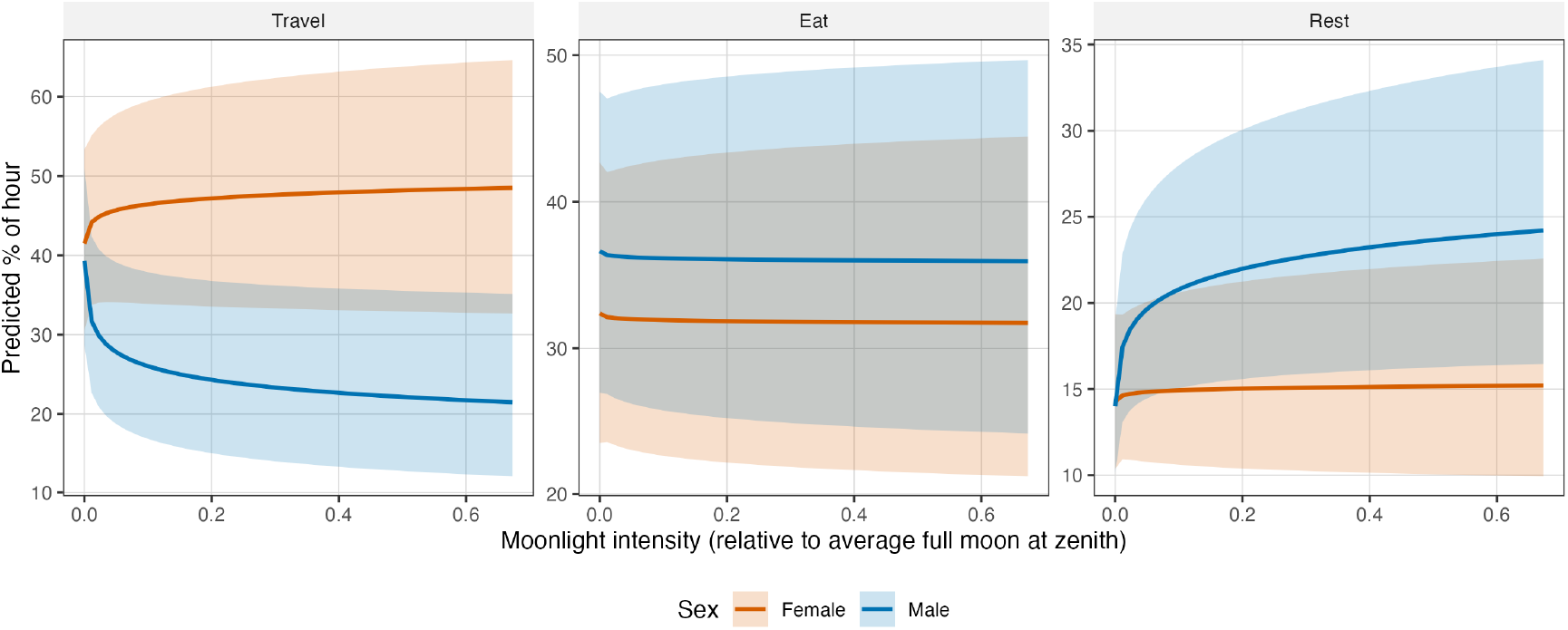
Predicted activity budget (percent time traveling, feeding, and resting per hour) as a function of moonlight intensity (clear-sky relative moonlight, scaled to an average full moon directly overhead), separately for females and males. Lines and bands as in Fig. 1.Moonlight had no significant main effect on any of the three activity-budget measures (all *P* > 0.4). The sex-by-moonlight interaction on percent time traveling was significant (*P* = 0.020); the parallel interaction on percent time resting was significant in the cloud-adjusted model (*P* = 0.040) but only marginally non-significant in the clear-sky model shown here (*P* = 0.093); the percent-time-feeding model did not include a sex-by-moonlight interaction term.

An exploratory secondary analysis of the probability that any social behavior was recorded in an hour also showed a significant sex-by-moonlight interaction (*P* = 0.028): the predicted probability of social behavior decreased with moonlight for females (from approximately 13% to 3% of hours) but increased with moonlight for males (from approximately 18% to 29% of hours) – the opposite pattern from the percent-time-traveling result above.

### Nightly Travel Distance

Fruit abundance had a significant, positive effect on nightly travel distance in the clear-sky model (β = 0.236, SE = 0.120, 95% CI [0.001, 0.470], *P* = 0.049) and a similar, marginally non-significant effect in the cloud-adjusted model (β = 0.232, SE = 0.119, 95% CI [-0.002, 0.466], *P* = 0.052), consistent with the activity-budget results above.

Moonlight had a significant, positive main effect on nightly travel distance in the clear-sky model (β = 0.196, SE = 0.097, 95% CI [0.005, 0.386], *P* = 0.044), and a positive but non-significant effect in the cloud-adjusted model (β = 0.150, SE = 0.083, 95% CI [-0.013, 0.313], *P* = 0.072). This positive average effect, however, masked a striking divergence between the sexes: the sex-by-moonlight interaction was marginally non-significant in the clear-sky model (*P* = 0.050) but became statistically significant in the cloud-adjusted model (*P* = 0.016). As moonlight increased, predicted nightly travel distance rose for females, from approximately 1,000 m to roughly 1,700-1,750 m, while it fell for males, from approximately 900 m to roughly 400-800 m (Fig. 3). This sex reversal was present under clear-sky moonlight but became more pronounced under the cloud-adjusted, realized-illumination version of the covariate, over which the male and female predicted curves diverge more sharply and across a narrower range of moonlight values (Fig. 3).

**Figure 3.**
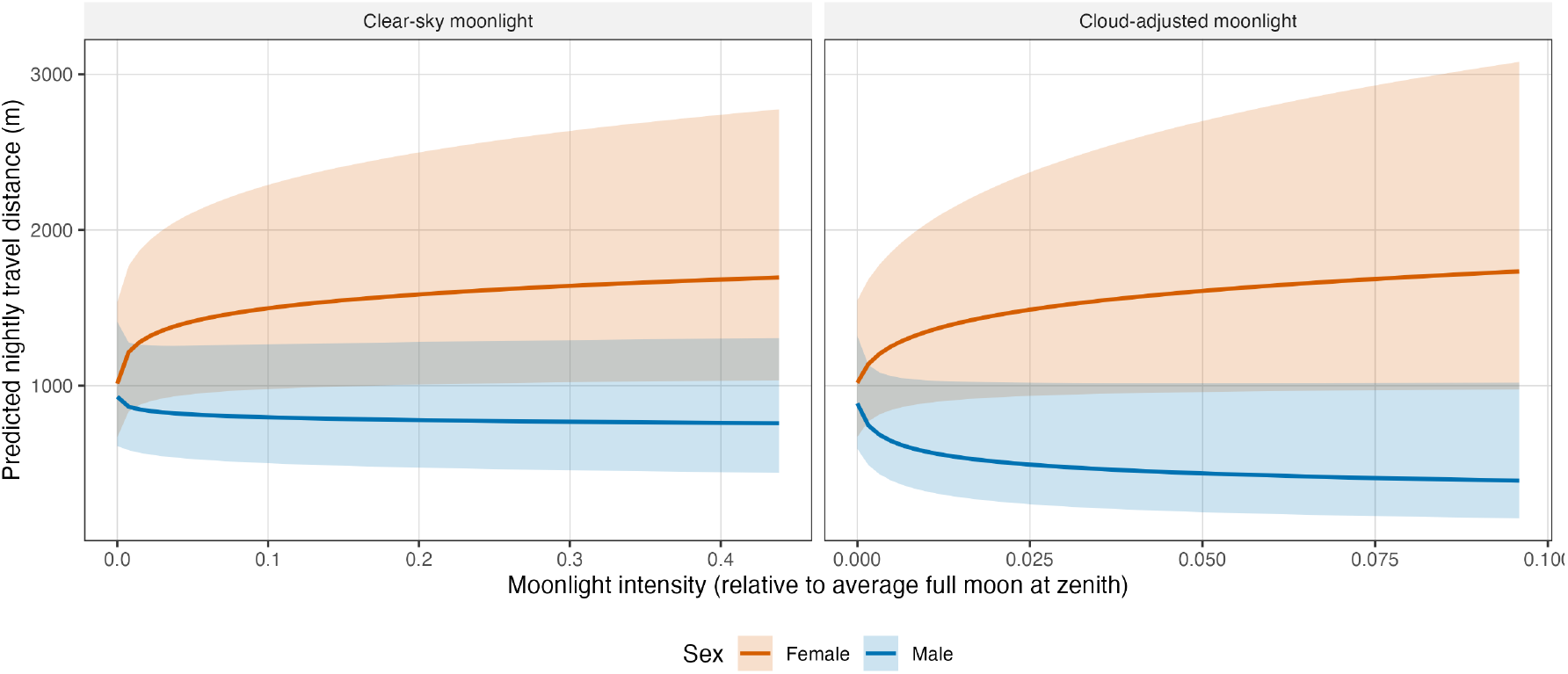
Predicted nightly travel distance as a function of moonlight intensity, separately for females and males, under the clear-sky moonlight index (left) and the cloud-adjusted, realized-illumination index (right). Lines are back-transformed predicted values from the linear mixed model of log nightly travel distance; shaded bands are 95% confidence intervals. Moonlight had a significant positive main effect on travel distance in the clear-sky model (β = 0.196, *P* = 0.044) and a non-significant positive effect in the cloud-adjusted model (*P* = 0.072). The sex-by-moonlight interaction was marginally non-significant in the clear-sky model (*P* = 0.050) but significant in the cloud-adjusted model (*P* = 0.016): predicted travel distance rose with moonlight for females (∼1,000 m to ∼1,700-1,750 m) and fell for males (∼900 m to ∼400-800 m), with the divergence sharper under the cloud-adjusted index. Because the two panels are built from the nightly travel-distance dataset, their moonlight axes span the nightly-mean range realized in that dataset (clear-sky maximum = 0.439; cloud-adjusted maximum = 0.096), which is narrower than the hourly range shown in Fig. 2.

### Rainfall

Rainfall showed no significant, consistent effect on any activity-budget proportion or on nightly travel distance once fruit abundance and moonlight were included in the models (percent time traveling: *P* = 0.267-0.274; percent time feeding: *P* = 0.058-0.063; percent time resting: *P* = 0.078-0.089; travel distance: *P* = 0.542-0.660; Fig. 4).

**Figure 4.**
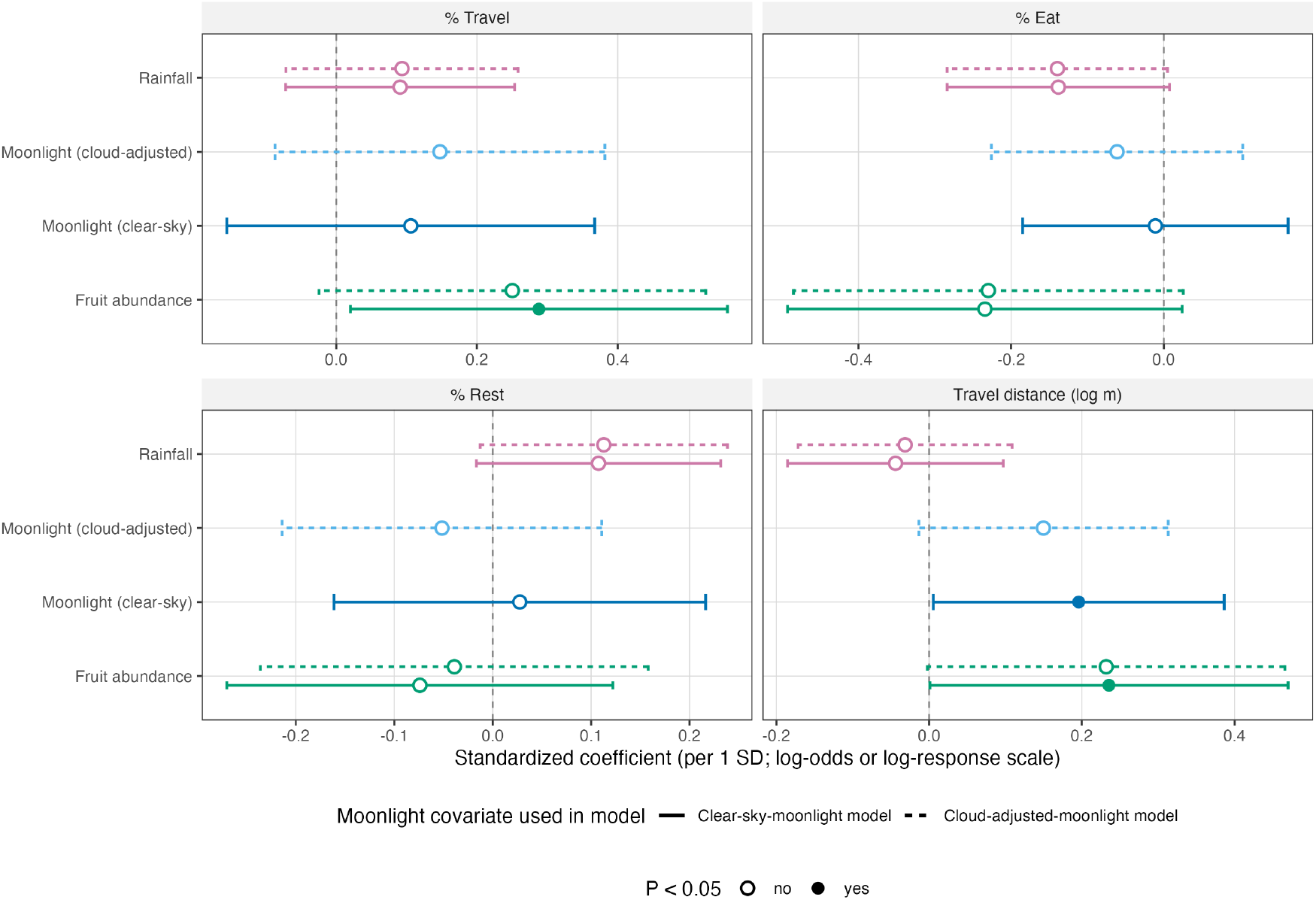
Standardized fixed-effect coefficients (points) and 95% confidence intervals (error bars) for fruit abundance, moonlight, and rainfall across the four primary response variables (percent time traveling, feeding, and resting; log nightly travel distance). Because moonlight was fit as two alternative covariates (clear-sky and cloud-adjusted) in otherwise identical model pairs, every other fixed effect shown here – including fruit abundance (green) and rainfall (pink) – was also estimated twice, once per model version; solid lines/points mark the clear-sky-moonlight model and dashed lines/points mark the cloud-adjusted-moonlight model, as indicated in the legend at the bottom of the figure. Filled points indicate *P* < 0.05. Fruit abundance was the most consistent predictor across response variables, with a significant positive effect on percent time traveling (clear-sky model, *P* = 0.035) and on travel distance (clear-sky model, *P* = 0.049).

### Model Comparison

Across all fitted response variables, the model summary and effect-size comparison (Fig. 4) show that fruit abundance was the most consistent predictor of activity allocation and movement, moonlight’s explanatory power was concentrated in its interaction with sex rather than in a uniform main effect, and rainfall added little independent explanatory value once fruit abundance and moonlight were accounted for. Residual diagnostics for the primary percent-time-traveling model showed no evidence of overdispersion (dispersion test *P* = 0.54) or outliers (outlier test *P* = 0.41), supporting the adequacy of the fitted beta GLMM for the reported activity-budget effects.

## DISCUSSION

Fruit abundance predicted both how kinkajous allocated their nocturnal activity and how far they traveled at night, but not in the direction we predicted: as fruit became more abundant, kinkajous spent a larger share of the night traveling and a smaller share feeding, and their nightly travel distances increased. We had expected the opposite, that scarcity would force longer search commutes between depleted patches. The pattern is instead consistent with kinkajous tracking and exploiting a rich resource landscape. In this forest, fruit-abundant periods are also periods when fruiting trees are closely spaced: the fruit-abundance index and the mean inter-fruiting-tree distance were strongly inversely related (r = −0.48), with fruiting trees averaging roughly 300 m apart during the most abundant censuses versus roughly 620 m apart during the sparsest. When many productive trees are available within a short distance of one another, an animal can profitably move among them, taking brief bouts at each, and the cost of doing so is low; when fruit is scarce and widely scattered, the same animal is better served by settling into longer feeding bouts at the few trees that are producing. Because kinkajous defend feeding areas around large fruiting trees against conspecifics (Kays 1999), high-abundance periods may also mean more encounters at more contested trees, further favoring movement over extended residence at any one patch. This is a resource-tracking rather than a search-cost account of the same trade-off between food acquisition and the costs of moving to get it (Terborgh and Janson 1986; van Schaik 1989).

The moonlight results reveal another unexpected pattern: male kinkajous traveled less as moonlight increased, while female kinkajous traveled more. The reduced travel under brighter moonlight seen in males is consistent with the classic predation-risk-avoidance response reported for many nocturnal mammals, in which increased conspicuousness under illumination favors reduced movement and increased vigilance or resting. The increased travel seen in females is the opposite, and is consistent with a movement-facilitation response, in which increased ambient light allows more, not less, nocturnal travel.

We motivated this movement-facilitation hypothesis at the outset by kinkajou body size and locomotor mode. As an animal larger-bodied than most of the nocturnal primates with which it shares the forest canopy, and dependent on continuous four-limbed contact with branches rather than confident leaping, a kinkajou risks real physical injury from a fall. If darkness itself constrains safe, efficient movement through the canopy for an animal with this locomotor style, then moonlight should facilitate, rather than suppress, travel and our results show that this is exactly what happened for females. We suggest that the sex reversal arises because the two sexes face a different balance between the predation-risk cost and the movement-facilitation benefit of moonlight. Being more food-limited by the costs of reproduction, females may have more to gain, in the currency of increased foraging range and efficiency, from traveling under brighter, safer illumination, even if this comes at some cost of increased visibility to predators. Males, without these reproductive costs, may instead be freer to prioritize the predation-risk-avoidance side of the trade-off, curtailing travel when moonlight is bright. This is the same reproductive-cost framework we invoke for the fruit results above, and it offers a single, consistent explanation for sex differences in both the feeding-competition and moonlight responses reported here.

The fact that this divergence became sharper and statistically significant under the cloud-adjusted, realized-illumination version of moonlight, while remaining only marginally non-significant under the clear-sky, theoretical version, supports a light-dependent behavioral mechanism rather than a spurious correlation with lunar phase or season: the sexes’ responses tracked the moonlight that actually reached the ground more tightly than the moonlight that would have reached the ground under a clear sky. This lends confidence that the underlying driver is illumination itself, and not some other seasonal or lunar-cycle covariate for which clear-sky moonlight might otherwise serve as an unintended proxy.

Kinkajous are frequently noted to converge ecologically with arboreal primates in their frugivory, prehensile-tailed climbing locomotion, and social structure (Kays and Gittleman 2001; Kays 2003). Our moonlight results suggest that this convergence has limits set less by absolute body size than by locomotor mode: some sympatric nocturnal primates are comparably sized to kinkajous, but where nocturnal primates typically move by leaping and can leap confidently between supports, the kinkajou is dependent on continuous four-limbed contact with branches and a prehensile tail. Consistent with a real locomotor cost to moving through the canopy in darkness, recent high-resolution tracking data show that kinkajous repeatedly use the same movement paths when climbing between trees rather than choosing new routes opportunistically each night (Gunner et al. 2026), suggesting that canopy movement in this species is spatially constrained and habitual in a way that would plausibly make illumination of the substrate valuable. A moonlight response that, at least for females, runs in the opposite direction from the lunar-phobic pattern typical of leaping nocturnal primates is consistent with this locomotor-cost framing.

Some caveats apply. Our behavioral sample comes from nine individually followed animals over one field season at a single site, so the generality of the sex-reversed moonlight response to other kinkajou populations, or to other years with different fruiting phenology, remains to be tested. The unequal number of half-nights per individual, typical of long-term focal-animal studies of a wide-ranging nocturnal frugivore, means that some individuals contribute more strongly than others to the sex-level patterns reported here; the individual random intercepts included in every model account for this repeated-measures structure but cannot fully substitute for a larger, more balanced sample of animals. Despite these limitations, the consistency of the female reproductive-cost framework across both the fruit-abundance and moonlight results, and the sharpening of the moonlight-sex interaction under the more biologically realistic cloud-adjusted illumination measure, together support a coherent account in which food limitation and locomotor constraints jointly shape how male and female kinkajous use moonlight and respond to fruit scarcity.

## ACKNOWLEDGMENTS

We thank the Smithsonian Tropical Research Institute for logistical support in Panama and the Instituto Nacional de Recursos Naturales Renovables (INRENARE) and other Panamanian park authorities for permission to conduct research in Parque Nacional Soberanía. We are grateful to R. Azipure, C. Carassco, C. Foster, N. Kays, C. Krieger, L. Slatton, D. Staden, and J. Young for their work following kinkajous in the field. J. Wright helped design the fruit census and A. Herre, O. Calderon, and E. Sierra helped identify fruits. D. Robinson and T. Robinson created and maintained the trail network at the Limbo research area that made this and companion studies possible. We also thank John Gittleman for advice on the evolutionary interpretation of these results and Matt Gompper for advice on the fieldwork. Claude (Anthropic) was used to assist with data cleaning, retrieval of environmental covariates, statistical model fitting, figure generation, and manuscript drafting; the author verified all analyses, results, and interpretations, and is solely responsible for the content.

## LITERATURE CITED

Brooks, M. E., K. Kristensen, K. J. van Benthem, A. Magnusson, C. W. Berg, A. Nielsen, H. J. Skaug, M. Machler, and B. M. Bolker. 2017. glmmTMB balances speed and flexibility among packages for zero-inflated generalized linear mixed modeling. The R Journal 9:378–400.

Chapman, C. A., L. J. Chapman, R. Wangham, K. Hunt, D. Gebo, and L. Gardner. 1992. Estimators of fruit abundance of tropical trees. Biotropica 24:527–531.

Clutton-Brock, T. H. 1991. The Evolution of Parental Care. Princeton University Press, Princeton, New Jersey.

Gunner, R. M., et al. 2026. High resolution data reveal fundamental steps and turns in animal movements. Ecological Monographs 96:e70069.

Hartig, F. 2022. DHARMa: residual diagnostics for hierarchical (multi-level/mixed) regression models. R package version 0.4.6.

Hersbach, H., B. Bell, P. Berrisford, S. Hirahara, A. Horanyi, J. Munoz-Sabater, J. Nicolas, C. Peubey, R. Radu, D. Schepers, A. Simmons, C. Soci, S. Abdalla, X. Abellan, G. Balsamo, P. Bechtold, G. Biavati, J. Bidlot, M. Bonavita, et al. 2020. The ERA5 global reanalysis. Quarterly Journal of the Royal Meteorological Society 146:1999–2049.

Kays, R. W. 1999. Food preferences of kinkajous (Potos flavus): a frugivorous carnivore. Journal of Mammalogy 80:589–599.

Kays, R. W. 2003. Social polyandry and promiscuous mating in a primate-like carnivore: the kinkajou (Potos flavus). Pp. 125-137 in Monogamy: Mating Strategies and Partnerships in Birds, Humans and Other Mammals (U. H. Reichard and C. Boesch, eds.). Cambridge University Press, Cambridge, United Kingdom.

Kays, R. W., and J. L. Gittleman. 2001. The social organization of the kinkajou Potos flavus (Procyonidae). Journal of Zoology 253:491–504.

Kays, R. W., J. L. Gittleman, and R. K. Wayne. 2000. Microsatellite analysis of kinkajou social organization. Molecular Ecology 9:743–751.

Milton, K. 1980. The Foraging Strategies of Howler Monkeys: A Study in Primate Economics. Columbia University Press, New York.

Palmer, M. S., A. Fieberg, A. Swanson, M. Kosmala, and C. Packer. 2017. A ‘dynamic’ landscape of fear: prey responses to spatiotemporal variations in predation risk across the lunar cycle. Ecology Letters 20:1364–1373.

Prugh, L. R., and C. D. Golden. 2014. Does moonlight increase predation risk? Meta-analysis reveals divergent responses of nocturnal mammals to lunar cycles. Journal of Animal Ecology 83:504–514.

Smielak, M. K. 2023. Biologically meaningful moonlight measures and their application in ecological research. Behavioral Ecology and Sociobiology 77:21.

Terborgh, J., and C. H. Janson. 1986. The socioecology of primate groups. Annual Review of Ecology and Systematics 17:111–135.

van Schaik, C. P. 1989. The ecology of social relationships amongst female primates. Pp. 195-218 in Comparative Socioecology (V. Standen and R. A. Foley, eds.). Blackwell, Oxford.

Zippenfenig, P. 2023. Open-Meteo.com Weather API [Computer software]. Zenodo. 10.5281/zenodo.7970649

